# Developmental Effects of the Pesticide Imidacloprid on Zebrafish Body Length and Mortality

**DOI:** 10.1101/395327

**Authors:** Akshay Krishnan, Christin Clyburn, Patrick Newcombe

## Abstract

The pesticide imidacloprid, a Neonicotinoid, is widely used and commercially available. Neonicotinoids are antagonists at nicotinic acetylcholine receptors, which can cause neurobehavioral effects in developing organisms. While the effects of other insecticides such as Malathion and Fipronil on zebrafish *Danio rerio* development have been studied, few studies describe the morphological effects of imidacloprid during zebrafish development. To test the hypothesis that imidacloprid concentration positively correlates to increased mortality and decreased body length, we exposed zebrafish to imidacloprid for five days post fertilization. Body length and embryo mortality were recorded at 1, 2, 3, 4, and 5 days post fertilization in 0 ug/L, 100 ug/L, 1,000 ug/L, and 10,000 ug/L imidacloprid concentrations. These concentrations were chosen to mimic levels that could be reached in the environment especially soon after application. Our results demonstrate statistically increased embryo mortality, with a f crit value of 3.193 and a f value of 3.098 from an ANOVA test, and impaired body length with a dose dependent correlation to the concentration of imidacloprid and a P value of less than 0.01 from a T test. Given the pesticide’s high prevalence, future studies should be considered to determine if this effect may impact livestock and human development.

## Introduction

Pesticides are commonly used in commercial and even residential applications. As a result, there is increased exposure to these agents either directly from plants and vegetables, or indirectly from agricultural water runoff. This has created concern which reaches all the way down to the local municipal authorities.^1^

The most common classes of pesticides are Organophosphates and Neonicotinoids, although the proportion of the latter has increased use due to their presumed effect to target insect over vertebrate nicotinic cholinergic receptors.^2^ This same beneficial mechanism of action may have unintended consequences for humans exposed to such agents. Many of these effects are direct and exist at very low levels. For example, if ingested, one seed treated with imidacloprid, a prototypic and widely used neonicotinoid, can kill an adult bird.^3^ Other effects are seen in developmental biology. The organophosphate Malathion has been documented to impair morphological development due to the inhibition of acetylcholinesterase (AChE), an enzyme that breaks down the neurotransmitter acetylcholine (ACh) to prevent excessive postsynaptic neuron and muscle stimulation that allows for body development.^4^ Studies have associated pesticide exposure with an increased rate of birth defects both in farm workers as well as non-farm workers who live within close proximity to increased pesticide use.^5^

One model for studying these development effects is to use zebrafish (*Danio rerio*), a small fish in the family cyprinidae.^6^ Given the similarities in vertebrate development between these fish and humans combined with the rapid growth rate, these fish have been extensively used in the literature to study the effect of various agents on developmental biology.^7^ Prior studies have used this model to demonstrate the specific effect pesticides may have on congential defects.^8,9^ One study has shown that the pesticide Fipronil (a non neonicotinoid) causes problems in development of spinal locomotor pathways by blocking the glycine receptor in zebrafish, manifested by decreased body length, notochord degeneration, and abnormal axial muscle morphology.^10,11^ The effect of neonicotinoids, though, may be more severe in neurodevelopment biology given their mechanism of action as antagonists at nicotinic acetylcholine receptors.^12^ Neurobehavioral effects such as decreased larval swimming activity have been documented as a result of exposure to imidacloprid, and imidacloprid increases mortality rates in early development of the common carp.^2^

However, the potential detrimental effects of imidacloprid on body length and mortality have yet to be conclusively identified in zebrafish. This may be an important surrogate for the impact of imidacloprid on human development biology. We therefore sought to identify the morphological effect of escalating concentration exposure to imidacloprid during the 5 days post fertilization (dpf) on these two factors in zebrafish embryos.

## Materials and Methods

### Animals

Zebrafish embryos were bred from six fish (4 males and 2 females) in a breeding tank set up in the afternoon. The following morning, the tanks were checked for embryos. A total of 456 embryos were obtained and were used as described below.

### Imidacloprid Sourcing and Preparation

Compare-N-Save brand was obtained commercially from a local home improvement store. Imidacloprid was handled with the protection of safety goggles and gloves, and the bottle was shaken before use.

Concentrations were prepared by a measurement of .0012 g of imidacloprid (1.47% concentration) dissolved in 120 ml of previously prepared embryo water, resulting in a solution of 10,000 ug/L. This base solution was further diluted by powers of 10, using 10 ml of the higher concentration and 90 ml of regular embryo water to create concentrations of 0 ug/l(50 ml of embryo water as the control), 100 ug/L, 1000 ug and 10000 ug/L respectively. A control solution was prepared by measuring 50 ml of embryo water. The described dilution procedure was repeated daily to create constant exposure to the desired concentration with daily water changes for each of the 5 days. In the first trial, each imidacloprid dose was measured separately for each concentration, repeated each day to change the water.

### Exposure Protocol

We exposed 129, 129, 69 and 129 embryos respectively to concentrations of 10,000 ug/L, 1,000 ug/L, 100 ug/L and 0 ug/L(control) of imidacloprid. Embryos were transferred into these within at most 4.5 hours post fertilization, though earlier when time allowed. Only live embryos were transferred from breeding tanks to before exposure to imidacloprid. Exposure occurred in petri dishes, which were stored in an incubator at 28.5 degrees Celsius.

The first trial consisted of approximately 180 embryos that were divided into six petri dishes, two each of control, 1,000 ug/L, and 10,000 ug/L.

The second, third, and fourth trials each consisted of 120 embryos divided into four petri dishes of control, 100 ug/L, 1,000 ug/L, and 10,000 ug/L. In total, the four trials used 420 embryos.

### Imaging

Each day through 5 dpf, embryos were photographed under a compound light microscope after using the embryo swirl technique to group them into the center. Once the embryos became too active after hatching from their chorions, they were photographed without the aid of the dissectoscope. An iPhone 6 using iOS v 10 (Apple, Cupertino California) was used for all photographs. ImageJ (National Institutes of Health, Bethesda, MD), an imaging processing program was used to obtain calibrated embryo lengths.

## Results

[Figure 1] In the control, a rapid increase of body length occurred between 1 dpf and 3 dpf, while between 2 dpf and 4 dpf in 100 ug/L, 3 and 5 dpf in 1,000 ug/L, and not beginning until 4 dpf in 10,000 ug/L. The first substantially increased body length measurements occurred, respectively per concentration, at 2 dpf, 3 dpf, 4 dpf and 5 dpf, each concentration increase corresponding to another day of development. By 5 dpf, the body lengths moved closer to each other, though still showed a correlation between decreased body length and imidacloprid concentration. On average, control embryos were 0.05 mm longer than 100 ug/L embryos, 100 ug/L embryos 0.20 mm longer than 1,000 ug/L embryos, and 1,000 ug/L embryos 0.31 mm longer than 10,000 ug/L embryos for a percent difference of 98.6%, 92.8%, and 84.0% compared to the control, respective in the order of concentration.

**Figure 1:**
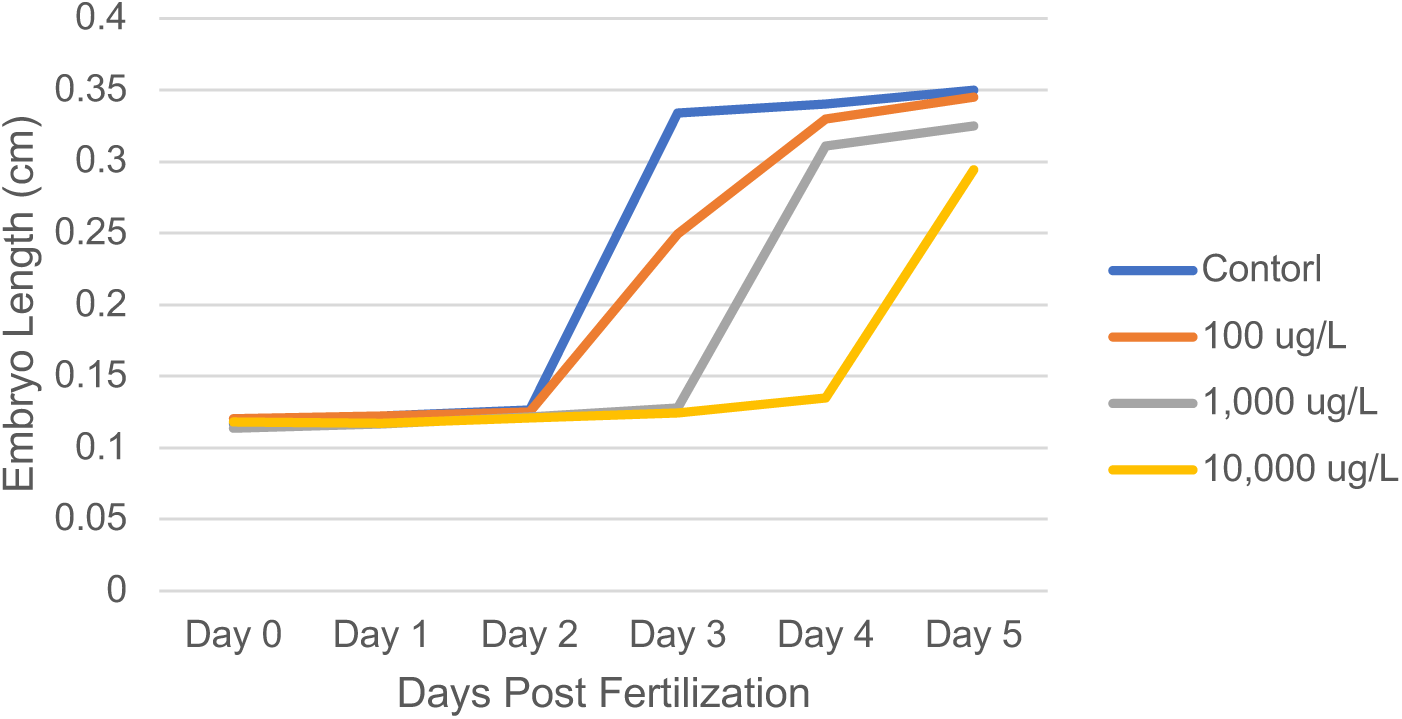
Embryo Lengths. This graph depicts the average embryo length from fertilization through five days post fertilization, from across four trials. The first trial did not include photographs taken the same day as fertilization, and trials two and three missed 3 dpf and 4 dpf respectively. We made one graph summarizing all the trials because, on days when we could not photograph the embryos, we excluded those days from the total data so that all are data was represented. Consequently, between all the trials we had data for each day to accurately portray the data.

Despite the first trial using individual imidacloprid measurements as opposed to dilutions, the body length data was generally consistent between each trial with the exception of the 10,000 ug/L solution during the third trial, which is approximately .02 mm greater than the similar other three trials at 5 dpf. A Linear Regression Statistical Test was performed and found that the data had significant statistical differences from days two through five, with the P values being consistently below 0.001 for 1000 and 10000 ug/L from day two to day five. However, the P values for the 100 ug/L were 0.04.

[Figure 2] Embryo mortality was also positively correlated to the concentration of imidacloprid. Through 1 dpf, the mortality rates in 100 ug/L and 1,000 ug/L were identical, and 100 ug/L embryos experienced a greater number of embryos dead, but control and 10,000 ug/L were always, respectively, above or below the other two. After 3 dpf, 1,000 ug/L demonstrated increased mortality than 100 ug/L. After 2 dpf, the distinction between 100 ug/L and 1,000 ug/L strengthened. By 5 dpf, the average number of living embryos ranged from 28 in control to 21 in 10,000 ug/L, a 23.33% difference. Control embryos experienced a 7.25% mortality rate, 100 ug/L embryos a 17.39% mortality rate, 1,000 ug/L embryos a 24.64% mortality rate, and 10,000 ug/L embryos a 33.33% mortality rate. Mortality rates were not recorded after 5 dpf because of a fungal infection that could not be adequately controlled.

**Figure 2:**
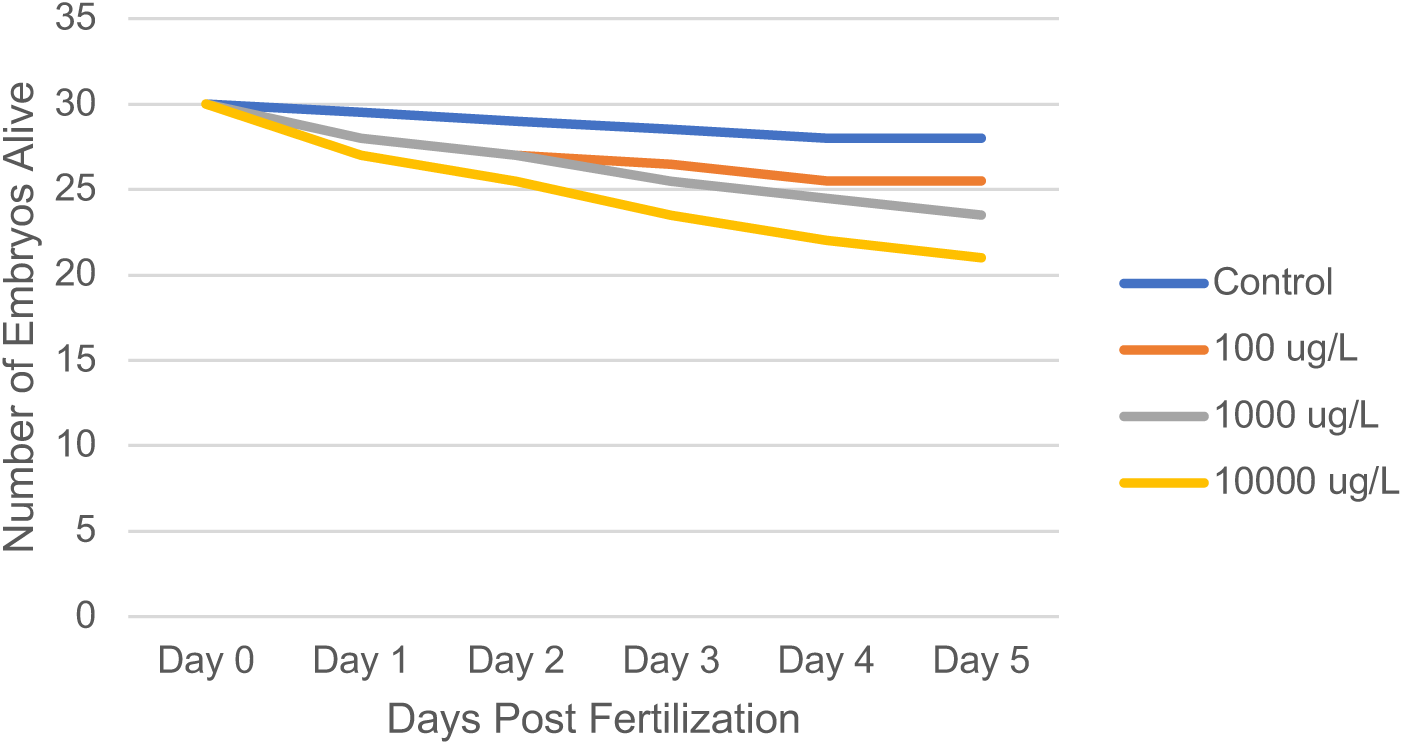
Mortality Rates. This graph shows the average mortality rate in each trial run at each day post fertilization. Embryo mortality was positively correlated to the presence of imidacloprid.

[Figure 3] Each power-of-ten increase in imidacloprid concentration correlated to between 5 and 10 more embryos to deaths, and out of the 276 embryos the chart draws from. Mortality at 5 dpf increased by 240% between control and 100 ug/L, more than doubling, increased by 142% between 100 ug/L and 1,000 ug/L, and increased by 135% between 1,000 ug/L and 10,000 ug/L, demonstrating the percent increase in embryo mortality to positively correlate to the smallest imidacloprid doses. An ANOVA statistical test resulted in statistically significant results that supported the hypothesis that imidacloprid concentration is positively correlated to embryo mortality. It produced a f crit value of 3.193 and a f value of 3.098

**Figure 3:**
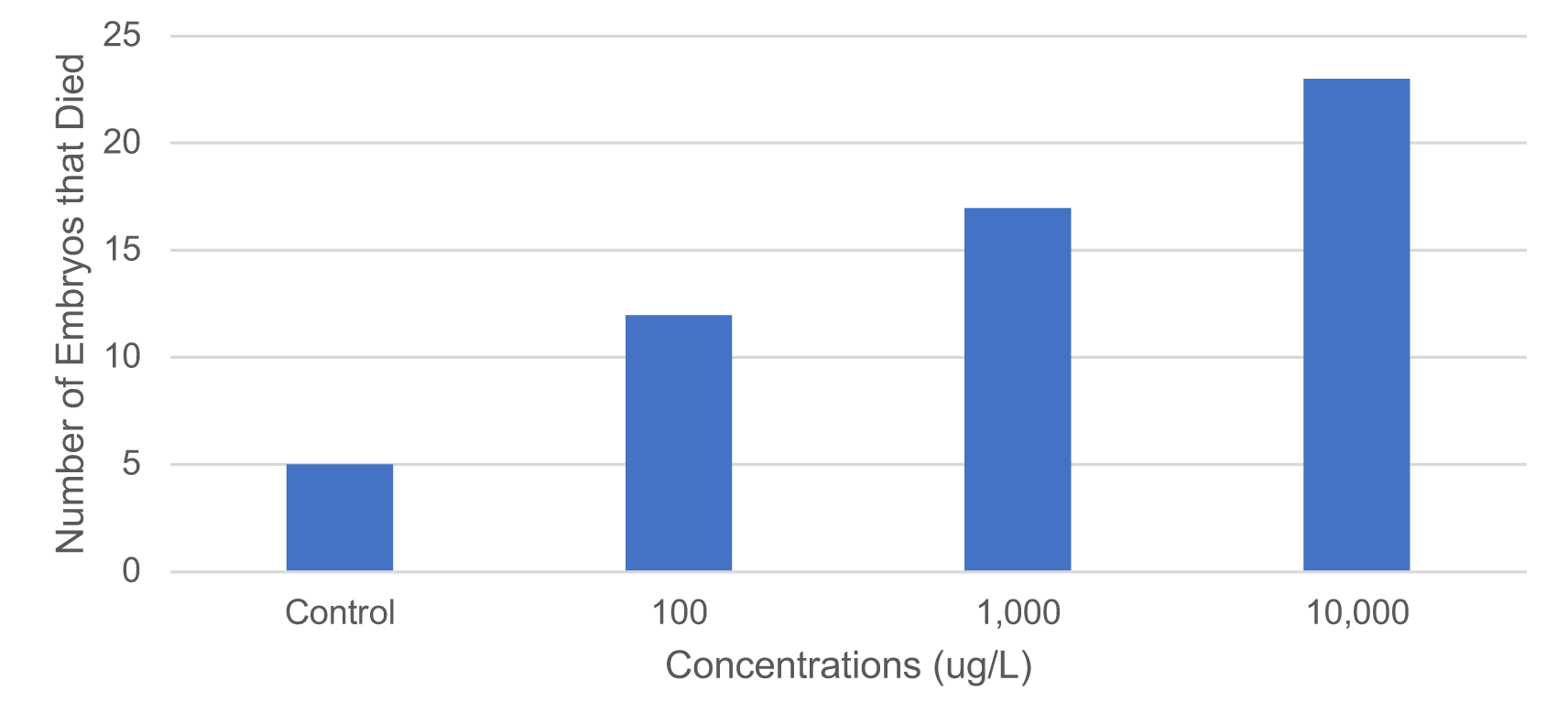
Total Mortality Per Concentration. This graph shows the total embryo mortality in the experiment across trials 2, 3, and 4 at 5 dpf. Each concentration held 69 embryos for a total of 276.

## Discussion

Pesticide exposure has been associated with neurodevelopment delays in various embryo models.^13^ However, limited data exists on these impacts for a common class of pesticides, neonicotinoids, on important developmental milestones. We used zebrafish as a model to determine the effect of pesticide exposure on two important parameters, embryo length and mortality. Our results demonstrated that imidacloprid impairs body length development in zebrafish embryos, in a dose dependent fashion confirming the hypothesis. More specifically, the rapid period of growth occurring in each concentration, likely the stage in which embryos abandoned their chorions, was delayed by approximately one day per power-of-ten increase in imidacloprid concentration as a result of exposure to imidacloprid, indicating that imidacloprid impairs body length development at 100 ug/L, 1,000 ug/L, and 10,000 ug/L.

As a measurement of growth, body length can represent both cellular and molecular changes.^4^ Imidacloprid’s action as an agonist at nicotinic acetylcholine receptors is found in mammals and insects, and these impacts may extend to zebrafish despite what was previously suggested.^2^ Organophosphates impair body length by breaking down the neurotransmitter acetylcholine (ACh) to prevent excessive postsynaptic neuron and muscle stimulation that allows for body development, thus leading to decreased body length in zebrafish and xenopus development.^4^ Conceivably, such a mechanism in zebrafish would elicit similar results if it, too, breaks down acetylcholine. Thus, given imidacloprid’s neurobehavioral effects on zebrafish larval swimming activity, novel tank exploration, and sensorimotor response, reduced body length may come from harm to the neurotransmitter acetylcholine as in organophosphates.^2,4^ Further research should address the longevity of impairments to body length development.

Our study also demonstrated that imidacloprid significantly increases mortality in zebrafish, suggesting that the neonicotinoid reduces the likelihood of embryo survival, and as a result could limit populations. Moreover, the greatest increase in embryo mortality between concentrations occurred between control and 100 ug/L, illustrating the significant impact of imidacloprid. This decline was observed at all of 100 ug/L, 1,000 ug/L, and 10,000 ug/L and supporting the dose dependent nature of the findings. A similar increase in mortality rates in the common carp has been described as a result of imidacloprid exposure.^11^ This same observation has been made for a sister pesticide, the organophosphate Malathion which has been shown to increased mortality during zebrafish development.^4^

The finding that the use of a common agricultural pesticide may impair body length development and result in increased mortality suggests that water runoff could harm aquatic vertebrate development in the environment. Various studies have documented the presence of pesticides in water systems around the world.^14–16^ This same agent has been implicated in developmental issues in other species including honey bees, and vetebrate life forms.^10,17^

In addition to the aquatic animal risk, our findings may suggest increased risks for humans exposed to this same pesticide. The zebrafish and human genomes are remarkably similar, sharing approximately 70% of their orthologues.^18^ Therefore, zebrafish are considered a very relevant model for human toxicology research. Studies have already shown an association between exposure to these agents and development defects like cleft palate and neural tube defects as well as congenital heart issues.^19,20^ More specifically correlated with our findings, studies in the literature have demonstrated low birth weight with chronic pesticide exposure as well.^21^

In summary, our findings demonstrated an effect on embryo length and mortality amongst Zebrafish. This species has been used as a model for other animals, suggesting similar findings may be seen in affected humans as well. Therefore, future study to address the impact of imidacloprid on livestock and humans should be considered.

## Acknowledgements

Sidwell Friends School biology instructor Melanie Fields offered guidance on the project. Director of Science and Agricultural Policy at the Chesapeake Bay Foundation Beth McGee generously offered her advice on our experiment, as did Washington College undergraduate Ernest Eckendrode, Davidson College Professor and Chair of Biology Dr. Barbara Lom, NIH chemist Dr. Heather Kalish, and NIH scientist Dr. Benjamin Feldman. Epidemiologist and Chief Medical Officer at Davita Clinical Research Dr. Steven Brunelli generously offered his skill in choosing and performing statistical tests.

